# Long-term memory in the migration movements of enucleated *Amoeba proteus*

**DOI:** 10.1101/125054

**Authors:** Carlos Bringas, Iker Malaina, Alberto Pérez-Samartín, María Dolores Boyano, María Fedetz, Gorka Pérez-Yarza, Jesus M. Cortes, Ildefonso M. De la Fuente

**Author notes:** Corresponding Author: Correspondence and requests for materials should be addressed to Ildefonso Mtz. de la Fuente. Instituto CEBAS-CSIC Campus Universitario de Espinardo. Espinardo. 30100 Murcia. ESPAÑA. /.

## Abstract

How motile, free unicellular organisms maximize the rate at which they encounter resources and develop optimal search strategies remains largely unknown. In fact, cell foraging is a very complex activity in which unicellular organisms integrate a diversity of external cues and develop efficient systemic movements to localize nourishment. These foraging strategies are critical when cells face conditions of scarce resources or they don’t possess information on where food is located. Here, in order to determine whether nuclear activity is directly involved in cell migration, we placed single, well-isolated, enucleated and non-enucleated starved *Amoeba proteus* on nutrient-free petri dishes, and we then analyzed their trajectories of movement using non-linear dynamic tools. We found that despite being enucleated, the systemic responses of the protoplasm exhibited typical biological behaviors, moving with apparent normality, creeping along the substrate, developing pseudopodia and gobbling up prey. Our quantitative studies show that both the non-enucleated and enucleated amoebas display a similar migration structure, characterized by super-diffusivity, non-trivial long-term correlations and move-step fluctuations with scale invariant properties. In conclusion, the nuclear activity does not seem to directly control the systemic cellular movements involved in locating sparse resources.

## Introduction

For a wide range of organisms, from bacteria to mammals, cell motility is an essential feature of systemic cell dynamics. In free cells, migration is necessary for critical activities like locating food and avoiding predators or adverse conditions in order to enhance their chances of survival. In multicellular organisms, cellular locomotion is required for a plethora of fundamental physiological processes, such as embryogenesis, tissue morphogenesis, organogenesis, tissue repair and immune responses. Indeed, deregulatedcellular migration is involved in important human diseases,^1, 2^ includingimmunodeficiencies and cancer’. In general terms, cellular life would be impossible without regulated motility. However, while some progress is being made in understanding this process, how cells move efficiently through diverse environments, and migrate in the absence of external cues, is an important unresolved issue in contemporary Biology.

Recent studies on chemotaxis have provided evidence that cell locomotion in response to dynamic soluble gradients exhibit long-term correlations^3^. Other research on cell migration in the presence of external molecular guidance have shown robust correlations between cell speed and persistence time(the time needed for a cell to change its motion direction)^4^. In the absence of external stimuli, cell migration has been also described as persistent random walking^5, 6^. In addition, several studies have observed that certain cell locomotion patterns are consistent with Lévy walks ^7, 8^, although some controversy remains regarding the accuracy of Lévy flight foraging^9^.

In order to determine whether nuclear activity is directly involved in cell migration, we have quantitatively analyzed the trajectories of 40 *Amoeba proteus* in the absence of external cues, 20 of which had been enucleated by micromanipulation.

Historically, it is known that enucleated amoebas can stay alive for long periods, up to 14 days^10^. In those experiments, the full viability of the enucleated organisms was verified by adding again the nucleus 12 days after enucleation; interestingly, some amoebas were capable to resume their complete physiological functions, including cellular division and developing stable cultures ^10^.

Note that the length of the *Amoeba proteus*’ cellular cycle greatly varies depending on the environment, but under controlled culture conditions, it is usually about 24 hours long^11^.

In our study, 40 amoebas were starved for one day at the beginning of the experiments, after which, half of the cells were enucleated and the organisms (including the enucleated amoebas, also calledcytoplasts) wereindividually placed on nutrient-free petri dishes. The motility of each cell was recorded using a digital camera attached to a stereo microscope, acquiring images every 2 seconds over a period between t_min_ = 130 min and t_max_ = 279 min (mean = 223 min). The digitized trajectories were analyzed in the form of time series *P*(*t*) using non-linear dynamic tools. Finally, after recording the movement of each cytoplast, the absence of a nucleus was verified by Hoechst staining and fluorescence microscopy (see Methods, Figure 1 illustrates the main experimental procedures).

**Figure 1.**
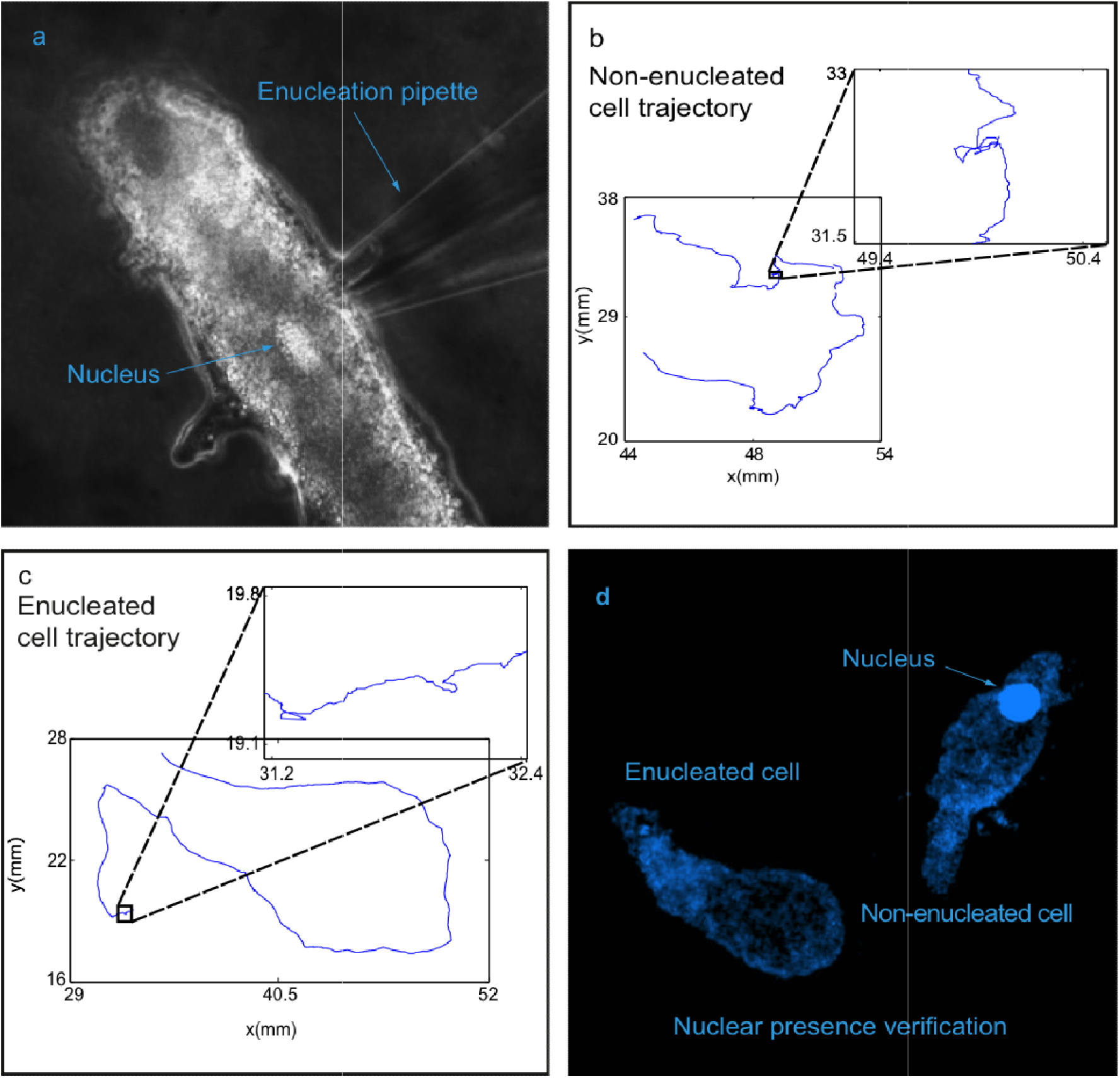
Experimental Procedures. *Amoeba proteus* cells were starved for 24 hours and half of them were enucleated using a micromanipulator: **a**, enucleation process. All the amoebas were placed in separate nutrient-free petri dishes, where their migration movements were recorded with a digital camera attached to a stereo microscope at 0.5 frames per second: **b** and **c**, digitized non*-*enucleated and enucleated cell trajectories, respectively. The inserts highlight local displacements by the amoeba. Finally, the presence or absence of a nucleus was verified in the enucleated amoebas. **d**, fluorescent microscopy image of an enucleated cell stained by Hoechst 33258 (1 mM) alongside a non-enucleated cell as a control.

## Results

The primary results showed that the enucleated amoebas behaved with apparentnormality in terms of adhesion to the substrate and overall motility. Moreover, in other experiments the cytoplasts efficiently hunted prey when exposed to it (see Supplementary video 1).On the other hand,it is also worth noting that in a preliminary observation all cell migrations were characterized by multiple short move-steps, which are occasionally alternated with long steps and stops (Supplementary video 2 and 3).

To quantitatively characterize the movements of the 40 organisms, we first analyzed the relative fluctuation in their trajectories (the deviation of the move-step length from its average) by computing the root mean square fluctuation (rms). This approach allows the existence of long correlations in the move-step fluctuations to be determined with precision (see Methods).

The net displacement in the move-step time series *u*(*t*)=*u*(1),*u*(2),…,*u*(*t*_*max*_) after *l* steps is defined as 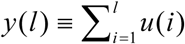, and the rms fluctuation of the average displacement is given by:

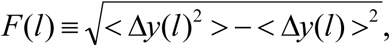

where Δ*y*(*l*) ≡ y(*l* +*l*_0_) – y(*l*_0_), and the brackets denote the average over all possible values of *lo.* Long-range correlations are detected if the fluctuations can be described by a power law such that *F*(*l*)*∼l*^*α*^. Thus, for uncorrelated data, the fluctuation exponent α is equal to 0.5. Markovian processes also give α = 0.5 for sufficiently large values of *l*, whereas α > 0.5 indicates the presence of positive long-range correlations and α < 0.5 implies long-term anti-correlations ^12^.

The *rms* analysis showed that(for R^2^ >0.989, coefficient of determination which measures the goodness fit)the α values of all the non-enucleated cells ranged between 0.645 and 0.9, with a global mean of 0.764 ±0.067 (mean ±SD, Supplementary Table 1). These non-trivial correlations encompassed between 200 and 3,000 movesteps (mean=1,125 ±798), which corresponded to periods ranging from 6.6 to 100 minutes (mean = 37.5 ±26.6). The enucleated cells exhibitedα values ranging between 0.625 and 0. 905, with a mean of 0.785 ±0.075 (Supplementary Table 2), corresponding to between 200 and 3,100 move steps (mean = 1,365 ±1,032)or equivalently to periods of time ranging from 6.66 to 103.3 minutes (mean = 45.5 ±34.4). Figure 2, panels a and b, show some examples of the rms analysis. The p-value of the Wilcoxon rank-sum test suggested that there were nosignificant differences in terms of the long-range correlations between non-enucleated and enucleated cells, neither in the correlation exponents (p-value = 0.365), nor in the duration of the correlations (p-value = 0.625). Likewise, the Detrended Fluctuation Analysis (see Methods) confirmed the existence of long-term correlations and scale invariance (Supplementary Table 1 and 2).

**Figure 2.**
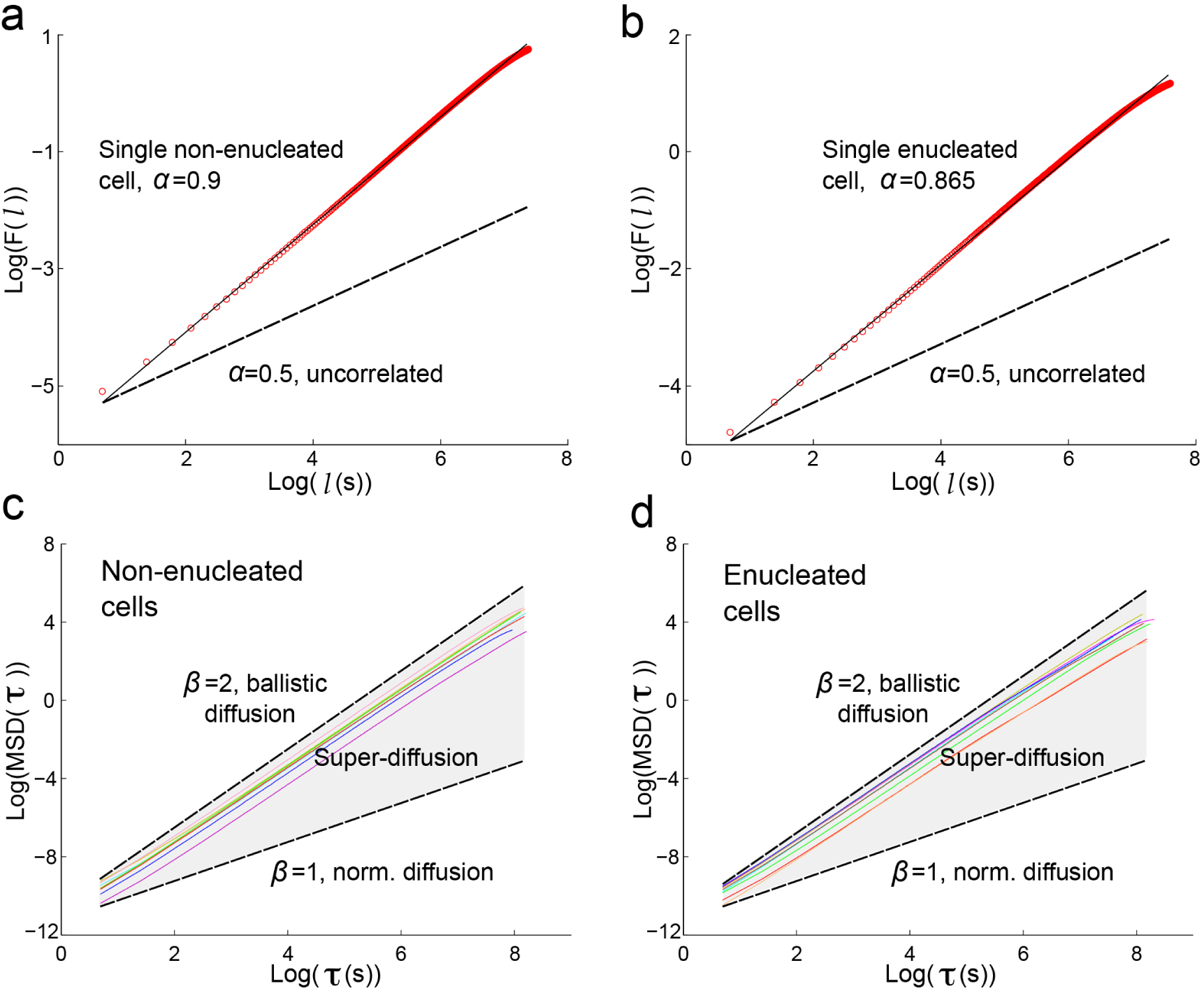
Root mean square fluctuation and Mean Square Displacement of the trajectories of non-enucleated and enucleated amoebas. Log-log plot of *rms* fluctuation *F* versus *l* step for a prototype non-enucleated cell (**a**), and a prototype enucleated cell (**b**). The slope for the non-enucleated cell was α=0.9, while for the enucleated it was α=0.865,indicating positive long-term correlations in both cases. In **c** and **d**, log-log plots of MSD against the time interval τ, for 8 prototypic non-enucleated and 8 enucleated cells, respectively. β=1 indicates normal diffusion while β=2 indicates ballistic diffusion. The grey region defines the area of super*-*diffusion, in which all experimental slopes are contained. The fact that τ_max_=1/4th of the data length, implies that super-diffusion holds in large scales.

Secondly, to quantify the amount of space explored by the amoebas during their locomotion, we calculated the Mean Square Displacement (MSD), which reflects the average square displacements over increasing time intervals between positions in a migration trajectory (see Methods). Specifically, this measure of the surface area explored by cells over time is related to the overall efficiency of migration^13^. As a relevant macroscopic property of random walks is known to involve the scaling that is related to the time taken to achieve the MSD according to the power law relationship *MSD*(τ)∼ Dτ^β^, where *β* characterizes the behavior of diffusive processes^13^. In a typical process, such as pure uncorrelated Brownian motion, the exponent *β* is equal to 1. If *β*>1 super-diffusion occurs, while if *β*<1 sub-diffusion takes place. Both processes are associated with a non-linear dependence of MSD over time, known as anomalous diffusion, which typically occurs in complex systems with long-range correlated phenomena. It should be noted that super-diffusion behavior has proven to be a more efficient foraging strategy under starvation conditions than diffusive searching^14,15^. Here, the MSD was calculated across both short and large timescales, and super*-*diffusion was observed in all cases, (Supplementary Tables 1 and 2; Figure 2, panels c and d, show an example of the MSD analysis for time windows as large as 1/4th of the amoeba data length). The *β* values obtained were compared with the Wilcoxon rank-sum test, which gave a p-value of 0.096, suggesting that there were no significant differences between the nourishment searching patterns of both groups.

Lastly, a global study of all the cell trajectories was performed by calculating the average speed and the directionality ratio. The average speed of the non-enucleated group was between 0.003 mm/s and 0.006 mm/s (mean = 0.004 ±0.001 mm/s) and it was not foundto be significantly different from the speed of the enucleated, which ranged from 0.002 mm/s to 0.006 mm/s (mean 0.004 = ±0.001 mm/s, p-value = 0.156 in the Wilcoxon rank-sum test: Supplementary Table 1 and 2). The directionality ratio quantifies the straightness of a trajectory^13^, where 1 represents a straight cell trajectory and approximating to 0 for a highly curved trajectory (see Methods). Again, no significant differences were found between the non-enucleated group, with a ratio comprehended between 0.082 and 0.717 (mean = 0.293 ±0.177) and the enucleated group, with ratios between 0.063 and 0.836 (mean = 0.332 ±0.185; p-value = 0.507 in the non-parametric test). Hence, the trajectories of both amoeboid groups were undistinguishable in their straightness.

## Discussion

In our study, we addressed quantitative aspects of *Amoeba proteus* migration in the absence of external stimuli, and the results show that the complex trajectories of both the enucleated and non-enucleated cells are well described by a particular kind of random motion characterized by super-diffusivity, non-trivial long-term correlations, and move-step fluctuations with scale invariant properties. In particular, we found long*-*term memory over a mean of 41.5 minutes, which corresponded to correlations lasting approximately 1,245 move-steps. Thus, the move-step dynamics in all organisms analyzed is strongly influenced by previous movements. These long-range correlations in spatial displacement provide a possible mechanism of critical functionality and they reflect long-term memory at the level of cell behavior. Inasmuch as there were no significant differences between the general characteristics of amoeba locomotion, both in enucleated and non-enucleated cells, we conclude that the nuclear activity does not seem to be directly involved in the migration of these organisms in the absence of external stimuli. This phenomenon is reported here for the first time.

On the other hand, super-diffusion was observed in all migration trajectories, both in short and large scales, which suggests that the efficiency of systemic movements to localize nourishmentwere preserved in amoebas after enucleation.

From a molecular point of view, the amoebas’ locomotion is controlled by complex metabolic networks, which operate as non-linear systems with dynamics far from equilibrium^16^. These biochemical networks involve an intricate interplay of multiple components of the cell migration machinery, including the actin cytoskeleton, ion channels, transporters, regulatory proteins such as the Arp2/3 complex or the ADF/cofilin family proteins^17 18^. As a consequence of the efficient self-regulatory activity of the metabolic networks, each amoeba seems to be endowed with the ability to orientate its movement toward specific goals in the external environment, thereby developing efficient foraging strategies even in conditions of sparse resources when there is limited or no information as to where food is located.

A large number of experimental studies on individual cells have demonstrated that important systemic behaviors are supported by astounding metabolic properties^16^. For instance, the single cellamoeboid *Physarumpolycephalum* can find the minimum-length solution between two points in a maze^19^, forming an optimized network closely approaching the purposely designed Tokyo railway system^20^ to resolve complex multiobjective foraging problems ^21^, and to learn and recall past events ^22^.

These and other advances in the study of the cellular systemic metabolic properties^23 24^ are bringing forth a new conceptual framework that defines the cell essentially as *adynamic metabolic reactorin* which most of the constituent biomolecules are synthesized and destroyed through cyclic biochemical reactions driven by enzymes. In turn, the enzymatic processes are integrated in functional networks that, on a global level, shape one of the most complex dynamic systems in nature^16^. As a consequence, millions of ions and molecules are continuously transformed with extreme precision through tightly regulated reactions, all of which are confined to a reduced space of microns. This extraordinarily well coordinated biochemical activity allows each cell to behave as anintegrated metabolic system, capable of self-organization and self-regulation, as well as adapting to its environment^16^.

In accordance to the aforementioned studies, the enucleated amoeba’s behaviors herein observed may be explained by the singular self-regulated properties of the cellular metabolic life.

Amoeboid organisms have survived for hundreds of millions of years, with the ability to hunt expeditiously^25^, a characteristic systemic behavior that does not seem to depend directly on the nuclear activity.

It is truly extraordinary that this dynamic metabolic reactor called cell, in which thousands upon thousands of simultaneous biochemical processes are being coordinated to synthesize the most complex molecules in the universe, can behave as an individual self-regulated metabolic entity capable of efficiently finding resources, adapt to the external environment and perpetuate itself in time.

## Methods

### Cell Cultures

*Amoeba proteus* (Carolina Biological Supply Company, Burlington, NC. Item # 131306) were grown at 21 °C on Simplified Chalkley’s Medium (NaCl, 1.4 mM; KCl, 0.026 mM; CaCl_2_, 0.01 mM), alongside *Chilomonaas* food organisms (Carolina Biological Supply CompanyItem # 131734) and baked wheat corns. At the beginning of the experiments the amoebas were starved for about 24 hours.

### Enucleation

Each amoeba was twice washed in Simplified Chalkley’s medium and after enucleated on standard petri dishes using a Sutter MP-235 micromanipulator. In short, a thin glass pipette was introduced in the amoeba’s cytoplasm through the cell membrane and the nucleus was then manually sucked out. It is necessary to state that the enucleation process is quite aggressive and causes an important injury on the cell’s membrane as well as extract a portion of the cytoplasm along the nucleus. Therefore, the technique was attempted a maximum of two times per cell. Once enucleated, the cytoplasts were left undisturbed for 15 minutes.

### Track recording, digitizing and significance

The enucleated and non-enucleated amoebas were individually placed on single nutrient-free petri dishes, where the motility of each cell was recorded using a digital camera attached to a SM-2T stereo microscope. The size of the observation field was of about 7×5 cm and images were acquired every 2 seconds, over a period between 130 (3,900 frames) and 279 (8,370 frames) minutes long, the median being 223 minutes (6,690 frames). These times depended on the amoeba staying within the field of vision and not moving outside of it; the criterion was to only consider the trajectories that lasted 2 hours or more straight inside. Since automated tracking software is often inaccurate^26^, we performed manual tracking using the TrackMate software in ImageJ(http://fiji.sc/TrackMate), as strongly suggested elsewhere^26^. Each track corresponds to an individual amoeba, a single cell was never recorded more than once. The significance of the results was established by means of the Wilcoxon rank-sum test because all normality hypotheses were rejected. 40 digitized trajectories, divided into two equally sized groups (enucleated and non-enucleated) were analyzed. This number is larger than what is considered sufficient (10 for each group) for the obtainment of significant data in biological studies^27^.

### Cell staining

After recording, the absence of a nucleus was verified by Hoechst staining and fluorescence microscopy. The cells were fixed in paraformaldehyde (4%) for 5 minutes and then permeabilized in Triton-X 100 (0.1%) for 5 minutes. Subsequently, the amoebas were stained with Hoechst 33258 (1mM) for 10 minutes and finally observed under an Olympus inverted fluorescence microscope (High Resolution and Analytic Microscopy, SGIker, UPV/EHU).

### Root Mean Square Fluctuation (rms)

The root mean square (rms) fluctuation is a classical approach in Statistical Mechanics, based on ideas already raised by Gibbs^28^ and Einstein ^29^, later developed and extensivelyutilizedto quantify physiological signals^30 31^.We applied the rms to assess whether there were long-term correlations in move-step fluctuations within a single time-series, in accordance with Viswanathan’s procedure ^12^. To calculate the rms and given a two-dimensional trajectory *P*(*t*)=(*x*(*t*), *y*(*t*)) with values equidistant in time (in our case the time series measured in millimeters and a value obtained every 2 seconds), the series of displacements *u*(*t*)*=u*(*1*)*,u*(*2*)*,…,u*(*t*_*max*_) is defined as:

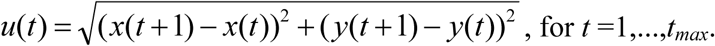

Then, the net displacement of the time series *u*(*t*) after *l* move-steps is computed as:

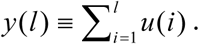

Finally, the rms fluctuation of the average displacement is calculated as:

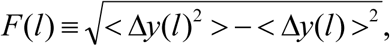

where Δ*y*(*l*) ≡*y*(*l +l*_0_) – y(*l*_0_), and the brackets denote the average over all possible values of *l*_0_. Thus, *F*(*l*) is defined as the square root of the difference between the average of the square of Δ*y*(*l*) minus the square of its average.

Long-range correlations can be detected if the fluctuations can be described by a power law so that *F*(*l*)∼*l*^α^. For uncorrelated data the fluctuation exponent α is equal to 0.5. Markovian processes also give an α = 0.5 for sufficiently large *l*, whereas α >0.5 indicates the presence of positive long-range correlations and α <0.5 implies long-term anti-correlations^12^. To calculate the number of correlated values, and therefore the duration of the correlation regimen, the value of *l* was systematically increased until the R^2^ adjustment decayed (implying a loss of the correlation).

### Mean Square Displacement

The MSD is a method proposed by Einstein in his work concerning Brownian motion, published in 1905^32^. Widely utilized since then (for example, to quantify bacteria motility^33^). This approach portrays the average squared displacements between positions in a migration trajectoryover increasing time intervals (or scales)^13^. Specifically, MSD can measure the surface area explored by cells over time and it is related to the overall efficiency of migration^14^.

Given a two-dimensional trajectory *P*(*t*)=(*x*(*t*), *y*(*t*)), *r*(*t*) is defined by:

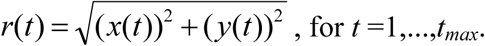

Thus, the MSD can be obtained by calculating:

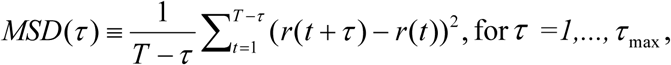

where *τ* denotes the time interval (or scale). Here, the diffusion was studied on both small and large scales, by setting the maximum time interval *τ*_max_ equal to 1/20th and to 1/4th of the data size respectively.

A relevant macroscopic property of random walks involves scaling in relation to the time of the MSD, with a power law relationship *MSD* (τ) ∼ Dτ^β^, where *β* characterizes the behavior of diffusive processes. In a typical process, such as pure uncorrelated Brownian motion, the exponent β is equal to 1. The phenomenon is called super-diffusion if 2>*β*>1, while if *β*<1 it is referred to as sub-diffusion. These two processes lead to a non-linear dependence of MSD over time, known as anomalous diffusion, which typically occurs in complex systems with long-range correlated phenomena.

### Detrended Fluctuation Analysis

Detrended Fluctuation Analysis (DFA) is a method proposed by Peng and coauthors in 1994^34^ to detect long-term correlations in time series that has been widely used to quantify physiological signals^35^. Given a time series *u*(*t*), we first obtain the signal profile by computing the cumulative sum of the series as:

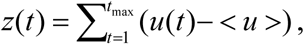

where the brackets denote the average of u(t). The time series obtained is then divided into boxes of equal length, *n* and the local trend *z*_n_(*t*) in each box is subtracted, the fluctuations of this detrended parameter and the integrated signal is calculated by:

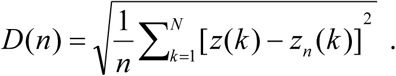

This computation is repeated for all box sizes, obtaining a relationship between fluctuations, *D*(*n*), and box sizes, *n*. A linear relationship on a log-log graph indicates the presence of scaling and under such conditions, fluctuations can be characterized by a scaling exponent γ. In general, the process is anti-correlated if 0< γ <0.5, while the process exhibits positive correlations when 0.5< γ <1.

### Directionality Ratio

The directionality ratio is a parameter that quantifies the straightness of a trajectory^13^, and it is equal to 1 for a straight cell trajectory, while it approaches 0 for a highly curved trajectory. Given a two-dimensional trajectory *P*(*t*)=(*x*(*t*), *y*(*t*)), we calculated the total length of the trajectory (*D*) by summing the series of displacements *u*(*t*):

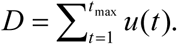

Next, the straight line (*d*) between the start point and the endpoint of the trajectory was calculated, and finally, the directionality ratio was computed as *d*/*D*.

## Acknowledgements

We would like to thank José Gonzâlez Romero and José Miguel Pérez Pérez from the Parasitology and Biomedicine Institute Lopez Neyra for their technical assistanceas well as the technical support provided by SGIker of UPV/EHU and European funding (ERDF and ESF).

## Author contributions

CB, IMDF: Designed experiments; CB, APS, MDB, GPY and MF: Performed the experiments; IM, JMC, IMDF: Performed quantitative analysis. All authors wrote the manuscript and agreed in its submission; IMDF: Conceived and directed the investigation.

## Data deposition statement

The datasets generated during and/or analyzed during the current study are available from the corresponding author on reasonable request

## Supplementary information

**Supplementary Video 1: Enucleated amoeba behavior.**

After being starved for 24 hours, a single amoeba was enucleated and placed individually in culture medium alongside food organisms. The enucleated cell presented apparently normal behavior, including the development of pseudopodia, adhesion to the substrate, cytoplasmatic streaming, exocytosis and phagocytosis, in this case of a single *Chilomona* cell followed by a fungal spore. In order to verify that the amoeba had been successfully enucleated, at the end of the experiment, the cell was subjected to fluorescent nuclear staining with Hoechst 33258 (1 mM), alongside a non-enucleated cell that served as a control (see Supplementary Methods 1).

**Supplementary Video 2: Digitized non-enucleated trajectory.**

The blue line represents the complete trajectory of the amoeba, the red circle represents the cell itself. This digitized video is an accurate representation of the cellular displacements sped up 60 times (from 13,238 seconds to 220).

**Supplementary Video 3: Digitized enucleated trajectory.**

The blue line represents the complete trajectory of the amoeba, the red circle represents the cell itself. This digitized video is an accurate representation of the cellular displacements sped up 60 times (from 13,642 seconds to 227).

**Supplementary Methods 1: Fluorescent staining corresponding to enucleated amoeba shown in Supplementary Video 1.**

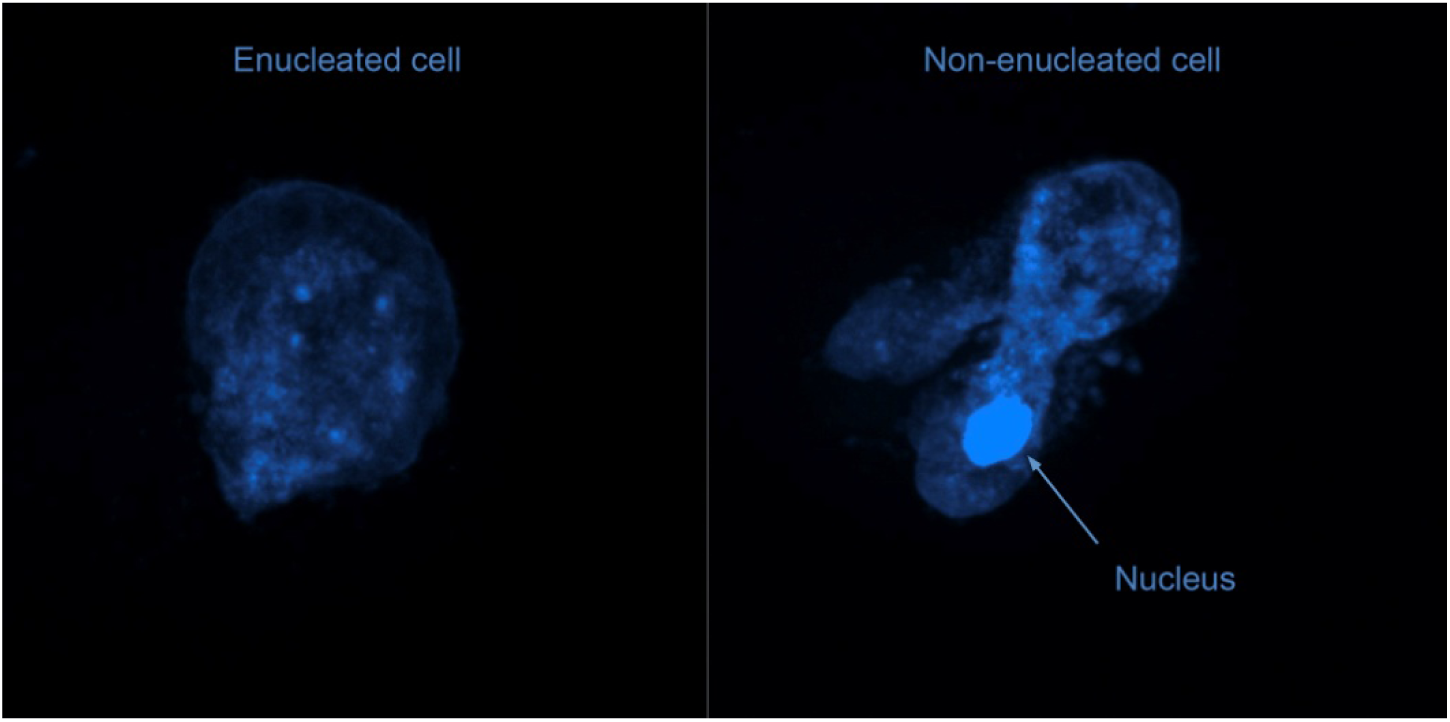

The enucleated amoeba recorded (see Supplementary Video1) was stained after recording with Hoechst 33258 (1 mM) alongside a non-enucleated control cell.

**Table S1.**
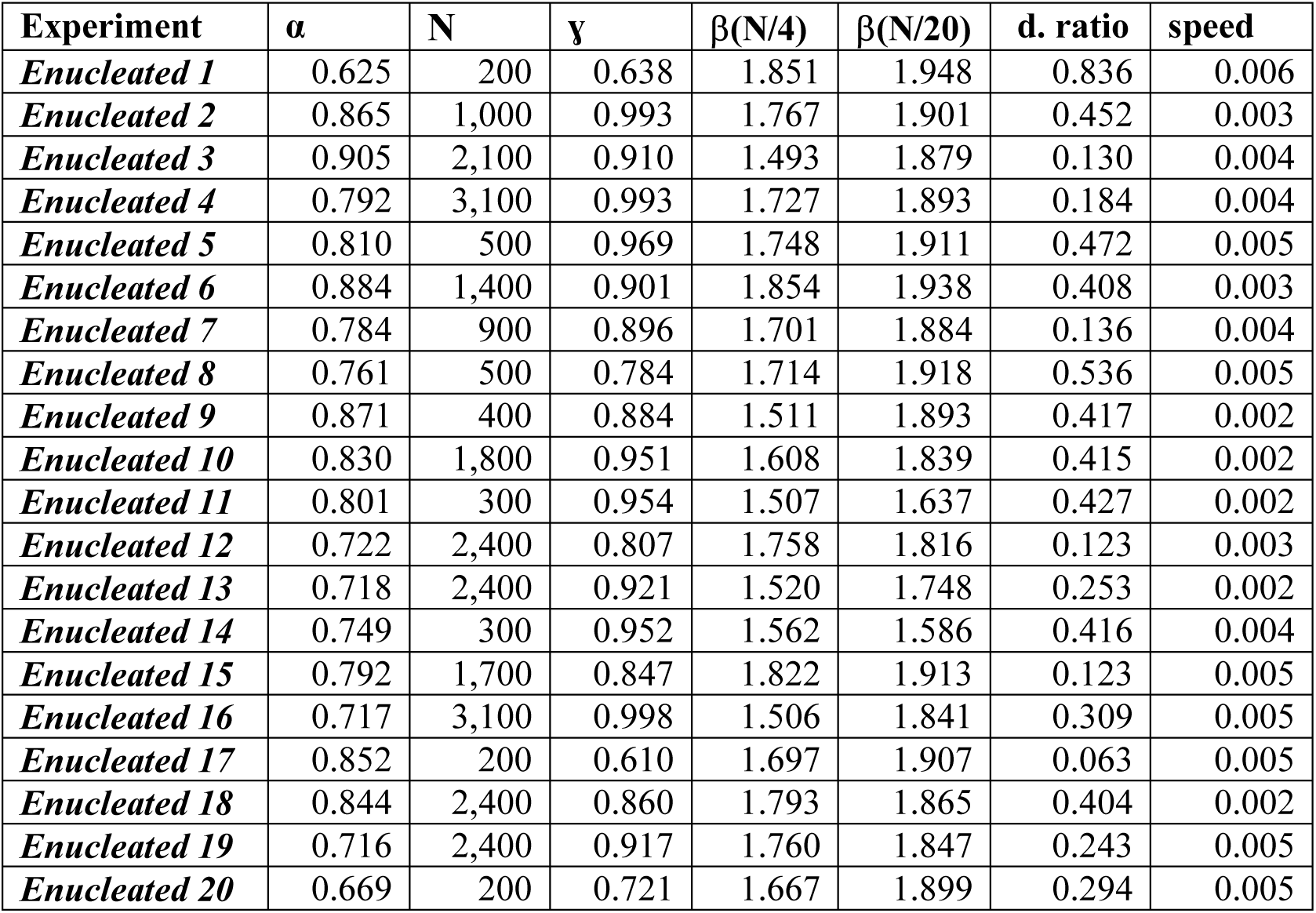
Quantitative statistics of enucleated amoebas.

The first column shows the number of the experiment, each one corresponding to a single non-enucleated amoeba. The rest of the data corresponds to the values ofrms correlation coefficient(α), number of move-length steps under the correlation regimen (C), DFA slope (γ), MSD slope for time intervals as large as 1/4th of the data size (β(N/4)),MSD slope for time intervals as large as 1/20th of the data size (β(N/20)), and directionality ratio, average speed measured in mm/s.

**Table S2.**
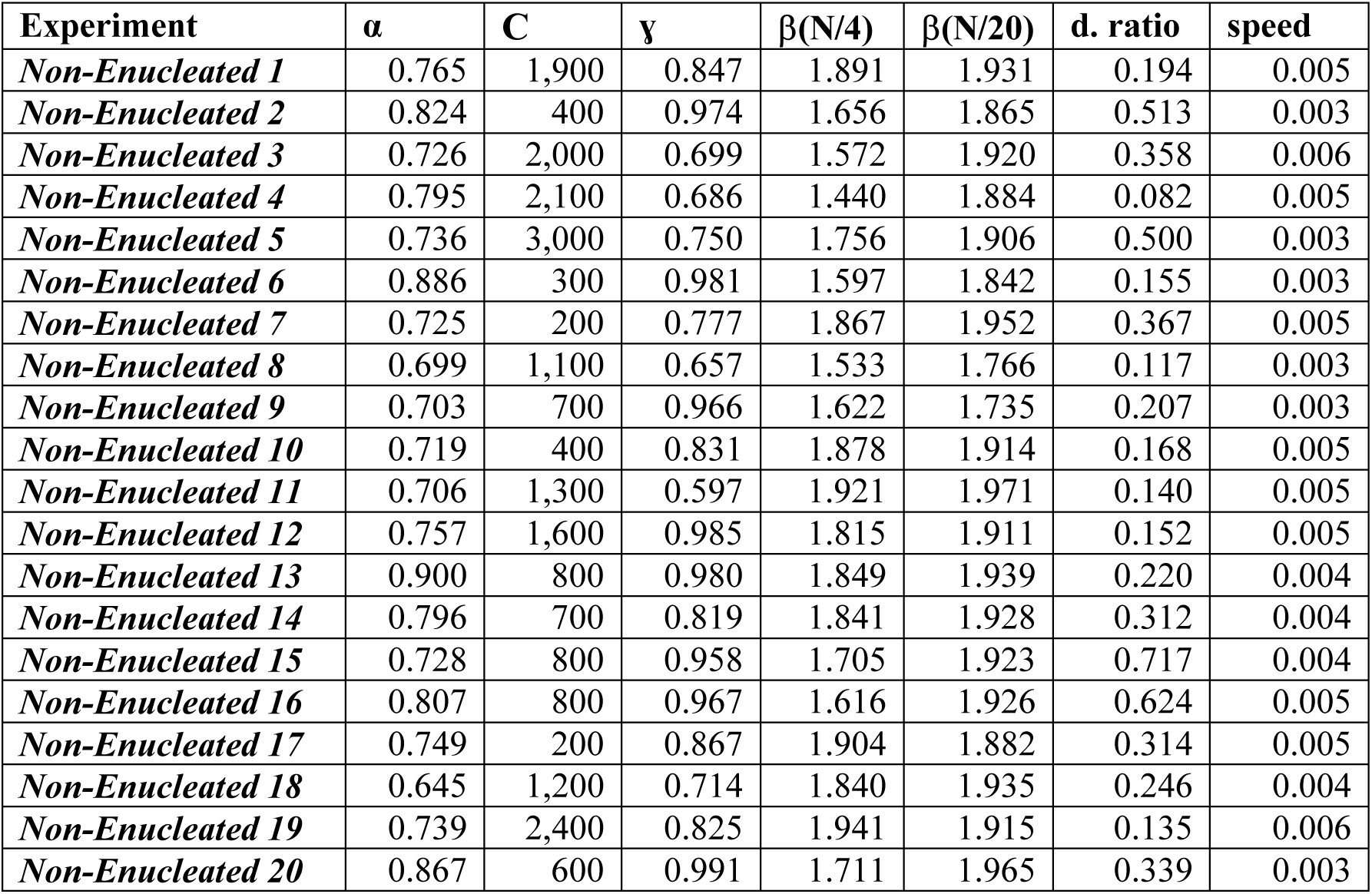
Quantitative statistics of non-enucleated amoebas.

The first column shows the number of the experiment, each one corresponding to a single enucleated amoeba. The rest of the data corresponds to the experimental values:rms, correlation coefficient (α), number of move-length steps under the correlation regimen (C), DFA slope (γ), MSD slope for time intervals as large as 1/4th of the data size (β(N/4)),MSD slope for time intervals as large as 1/20th of the data size (β(N/20)), directionality ratio, and average speed measured in mm/s.

## References

1. Bouma, G., Bums, S.O., & Thrasher, A.J. Wiskott-Aldrich Syndrome: Immunodeficiency resulting from defective cell migration and impaired immunostimulatory activation. Immunobiology. 214, 778–90 (2009)

2. Olson, M. F. & Sahai, E. The actin cytoskeleton in cancer cell motility. Clin. Exp. Metastasis. 26, 273–287 (2009)

3. Skoge, M. et al. Cellular memory in eukaryotic chemotaxis. Proc. Natl. Acad. Sci. USA. 111, 14448–53 (2014)

4. Maiuri, P. et al. Actin flows mediate a universal coupling between cell speed and cell persistence. Cell. 161, 374–86 (2015)

5. Selmeczi, D., Mosler, S., Hagedorn, P. H., Larsen, N. B. & Flyvbjerg, H. Cell motility as persistent random motion: theories from experiments. Biophys J. 89, 912–31 (2005)

6. Li, L., Cox, E. C. & Flyvbjerg, H. ’Dicty dynamics’: Dictyostelium motility as persistent random motion. Phys. Biol. 8, 046006 (2011)

7. Ariel, G. et al. Swarming bacteria migrate by Lévy Flight. Nat. Comm. 6, 8396 (2015)

8. Harris, T. H. et al. Generalized Lévy walks and the role of chemokines in migration of effector CD8+ T cells. Nature. 486, 545–8 (2012)

9. Pyke, G. H. Understanding movements of organisms: it’s time to abandon the Lévy foraging hypothesis. Meth. Ecol. Evol. 6, 1–16 (2015)

10. Ord, M.J. The viability of the anucleate cytoplasm of Amoeba proteus. J. Cell Sci. 1, 81–8 (1968)

11. Prescott, D.M. Relations between cell growth and cell division. I. Reduced weight, cell volume, protein content, and nuclear volume of amoeba proteus from division to division. Exp. Cell Res. 2, 328–37 (1955)

12. Viswanathan, G. M. et al. Lévy flight search patterns of wandering albatrosses. Nature. 381, 413–5 (1996)

13. Gorelik, R. & Gautreau, A. Quantitative and unbiased analysis of directional persistence in cell migration. Nat. Protoc. 9, 1931–43 (2014)

14. Viswanathan, G.M Raposo, E.P. & da Luz, M.G.E. Lévy flights and superdiffusion in the context of biological encounters and random searches. Physics of Life Reviews. 5, 133–150 (2008)

15. Faustino, C.L., da Silva, L.R., da Luz, M.G.E., Raposo, E. P. & Viswanathan, G. M. Search dynamics at the edge of extinction: Anomalous diffusion as a critical survival state. Europhysics Letters. 77, 30002 (2007)

16. De la Fuente, I. M. Elements of the cellular metabolic structure. Front. Mol. Biosci. 2, 16 (2015)

17. Artemenko, Y., Lampert, T. J. & Devreotes, P. N. Moving towards a paradigm: common mechanisms of chemotactic signaling in Dictyostelium and mammalian leukocytes. Cell. Mol. Life. Sci. 19, 3711–47 (2014)

18. Senoo, H., Cai, H., Wang, Y., Sesaki, H. & Iijima, M. The novel RacE-binding protein GflB sharpens Ras activity at the leading edge of migrating cells. Mol. Biol. Cell. 27, 1596–605 (2016)

19. Nakagaki, T., Yamada, H. & Tóth, A. Maze-solving by an amoeboid organism. Nature. 407, 470 (2000)

20. Tero, A. et al. Rules for biologically inspired adaptive network design. Science. 327, 439–42 (2010)

21. Dussutour, A., Latty, T., Beekman, M. & Simpson, S. J. (2010). Amoeboid organism solves complex nutritional challenges. Proc. Natl. Acad. Sci. U.S.A. 107, 4607–4611 (2010)

22. Saigusa, T., Tero, A., Nakagaki, T. & Kuramoto, Y. Amoebaeanticipateperiodicevents. Phys. Rev. Lett. 100, 018101 (2008)

23. De la Fuente, I. M. et al. Dynamic properties of calcium-activated chloride currents in Xenopuslaevis oocytes. Sci. Rep. 7, 41791 (2017)

24. De la Fuente, I.M., Cortes, J.M., Pelta, D.A., & Veguillas, J. Attractormetabolicnetworks. PLoSOne. 3, e58284 (2013)

25. Cosson, P. & Soldati, T. Eat, kill or die: when amoeba meets bacteria. Curr. Opin. Microbiol. 11, 271–6 (2008)

26. Hilsenbeck, O. et al. Software tools for single-cell tracking and quantification of cellular and molecular properties. Nat. Biotech. 34, 703–6 (2016)

27. Rosner, B. Fundamentals of Biostatistics. (Brooks/Cole, 2015)

28. Gibbs, J. W. Elementary Principles in Statistical Physics Developed with Especial Reference to The Rational Foundation of Thermodynamics. (Charles Scribner’s Sons, 1902)

29. Einstein, A. Zumgegenwärtigen stand des strahlungsproblems. PhysikalischeZeitschrift. 10, 185–93 (1909)

30. Ivanov, P. C. et al. Multifractality in human heartbeat dynamics. Nature. 399, 461–5 (1999)

31. Ivanov, P. C. et al. From 1/f noise to multifractal cascades in heartbeat dynamics. Chaos. 11, 641–52 (2001)

32. Einstein, A. Über die von der molekularkinetischenTheorie der WärmegeforderteBewegung von in ruhendenFlüssigkeitensuspendiertenTeilchen. Annalen der physik. 322, 549–60 (1905)

33. Long, Z. et al. Microfluidic chemostat for measuring single cell dynamics in bacteria. Lab Chip. 13, 947–54 (2013)

34. Peng, C. K. et al. Mosaic organization of DNA nucleotides. Phys. Rev. E. 49, 1685 (1994)

35. Goldberger, A. L. et al. Fractal dynamics in physiology: Alterations with disease and aging. Proc. Nat. Acad. Sci. U.S.A. 99, (suppl 1) 2466-72 (2002)

